# Can fMRI functional connectivity index dynamic neural communication?

**DOI:** 10.1101/2021.07.27.453965

**Authors:** Sonsoles Alonso Martínez, Alberto Llera, Gert ter Horst, Diego Vidaurre

**Author notes:** Department of Clinical Medicine, Center of Functionally Integrative Neuroscience, Aarhus University, Universitetsbyen 3, 8000 Aarhus, Denmark.

## Abstract

In order to continuously respond to a changing environment and support self-generating cognition and behaviour, neural communication must be highly flexible and dynamic at the same time than hierarchically organized. While whole-brain fMRI measures have revealed robust yet changing patterns of statistical dependencies between regions, it is not clear whether these statistical patterns —referred to as functional connectivity— can reflect dynamic large-scale communication in a way that is relevant to human cognition. For functional connectivity to reflect cognition, and therefore actual communication, we propose three necessary conditions: it must span sufficient temporal complexity to support the needs of cognition while still being highly organized so that the system behaves reliably; it must be able to adapt to the current behavioural context; it must exhibit fluctuations at timescales that are compatible with the timescales of cognition. To obtain reliable estimations of time-varying functional connectivity, we developed principal components of connectivity analysis (PCCA), an approach based on applying principal component analysis on multiple runs of a time-varying functional connectivity model. We use PCCA to show that functional connectivity follows low-yet multi-dimensional trajectories that can be reliably measured, and that these trajectories meet the aforementioned criteria. These analyses suggest that these trajectories might index certain aspects of communication between neural populations and support moment-to-moment cognition.

**Significance Statement:** fMRI functional connectivity is one of the most widely used metrics in neuroimaging research in both theoretical research and clinical applications. However, this work suffers from a lack of context because we still do not fully understand what fMRI functional connectivity can or cannot reflect biologically and behaviourally. In particular, can it reflect between-region neuronal communication? We develop methods to reliably quantify temporal trajectories of functional connectivity and investigate the nature of these trajectories across different experimental conditions. Using these methods, we demonstrate that functional connectivity exhibits reliable changes that are context-dependent, reflect cognitive complexity, and bear a relationship with cognitive abilities. These conditions show that fMRI functional connectivity could reflect changes in between-region communication above and beyond non-neural factors.

## Introduction

Communication between neuronal populations in the brain supports cognitive functioning. Like in any other complex system, communication in the brain can be understood as the inter-relationships of the various elements within the system, which are dynamically organised at many scales of space and time to generate complicated patterns of behaviour (Laughlin and Sejnowski, 2003). Here, we question whether functional connectivity (FC) between brain regions estimated from fMRI might at all reflect the communication system that subserves cognition.

The widespread existence of weak correlations between neurons in the brain is, in fact, a well-documented phenomenon (Averbeck et al., 2006; Averbeck and Lee, 2004; Cohen and Kohn, 2011; Nirenberg and Latham, 2003). An important question in theoretical neuroscience is about the mechanistic role of these correlations; particularly, whether or not these correlations convey information by themselves, i.e., as a separate “channel” beyond the firing rates and precise temporal ordering of the neuron’s firing events. If that were the case, then these correlations should be able to modulate their magnitude and configuration dynamically according to the external circumstances and the internal state of the animal (Cohen and Kohn, 2011; Nienborg and Cumming, 2009).

Here, we ask a similar question at the macroscopic level using human functional magnetic resonance imaging (fMRI) data, which, given its spatial coverage, might be better suited than the microscopic level to index higher-level aspects of cognition (Dehaene and Naccache, 2001). It is known that patterns of voxel activations can encode a variety of task-related variables (Hasson et al., 2009) and cognitive states (Haynes and Rees, 2006; Shine et al., 2019a). Furthermore, it has been shown that session-average patterns of covariance, or FC, between regions, are phylogenetically conserved (Lu et al., 2012), map the functional specialization of the different brain areas (Margulies et al., 2016), and can reflect clinical (Fox and Greicius, 2010) and psychological variations across the population (Finn et al., 2015; Karapanagiotidis et al., 2020; Smith et al., 2015). Our question of whether these (second-order) patterns of covariation can reflect actual large-scale communication is investigated through the assumption that cognition is subserved by large-scale neural communication, so we use cognition as a conceptual proxy between FC and communication.

There is strong evidence that the spatial patterns of FC exhibit change over time from seconds to minutes during both task execution (Cohen and D’Esposito, 2016; Shine et al., 2016; Xie et al., 2018) and periods of rest (Allen et al., 2014; Vidaurre et al., 2021; Vidaurre, 2021). These patterns of fluctuating FC understood as the temporal synchronization between signals across different brain areas can reveal important aspects of brain function (Allen et al., 2014; Sakoğlu et al., 2010; Zalesky et al., 2014). For example, it has been shown that FC reconfigures over time in response to internal and external circumstances (Chen et al., 2015; Elton and Gao, 2015; Hutchison and Morton, 2015) and that these reconfigurations may influence behaviour (Kucyi et al., 2017) and explain certain aspects of cognitive traits (Vidaurre et al., 2021). There is also evidence of moment-to-moment reconfigurations of FC in direct response to changes in environmental demands (Sadaghiani et al., 2015), indicating that fluctuations in FC can be relatively fast —relative to what fMRI can estimate, and that they can carry task-specific information. Capitalising on these findings, we propose three key aspects as necessary (yet not sufficient— see discussion below) conditions for FC to bear a meaningful relationship with neural communication: (i) these interactions should exhibit modulations of sufficiently high *temporal* complexity (since mental processes are dynamic and complex) at the same time than highly organised (so that the system behaves reliably); (ii) the nature of these modulations must be able to adapt to the behavioural context (i.e., responding according to external task demands or internal goals); and (iii) these modulations should span timescales compatible with the timescales of cognitive processes.

To find whether these conditions are met in real data, we developed a novel approach, principal components of connectivity analysis (PCCA), to provide reliable estimations of time-varying FC. PCCA combines multiple hidden Markov model (HMM) estimates from randomized initializations with principal component analysis (PCA) to extract the latent temporal structure across HMM runs (thereby accounting for the variability regarding model estimation and the complexity of the landscape of solutions). As opposed to previous approaches that made use of PCA (Shine et al., 2019b), PCCA captures the principal components, not of signal covariance, but changes in covariance —and therefore of changes in FC. In doing so, PCCA revealed a multidimensional axis of temporal covariation in macroscopic FC, which was found to fit all the above requirements. That is, we found robust trajectories of cross-region *covariation* that unfolded reliably across several dimensions (first condition), that were subject-specific and strongly modulated by the behavioural condition of the experiment (second condition) and did so at relatively short timescales in relation to the externally imposed cognitive demands (third condition). These results suggest that the estimated low-dimensional FC trajectories can not only be measured reliably but also suit key requirements to potentially index dynamic interregional communication.

## Results

We analysed fMRI data (repetition time, TR = 0.72s) from the Human Connectome Project (HCP; van Essen et al., 2013) from 100 subjects across three behavioural conditions: resting-state (rest), working memory (WM) task, and motor task. As illustrated in **Figure 1A**, we used the HCP resting-state data projected onto 25 independent components (ICs) as input time series, as well as task data obtained by applying the same (resting-state) group 25-IC spatial soft-parcellation on the HCP WM and motor task datasets (see *Methods* section for more details).

**Figure 1.**
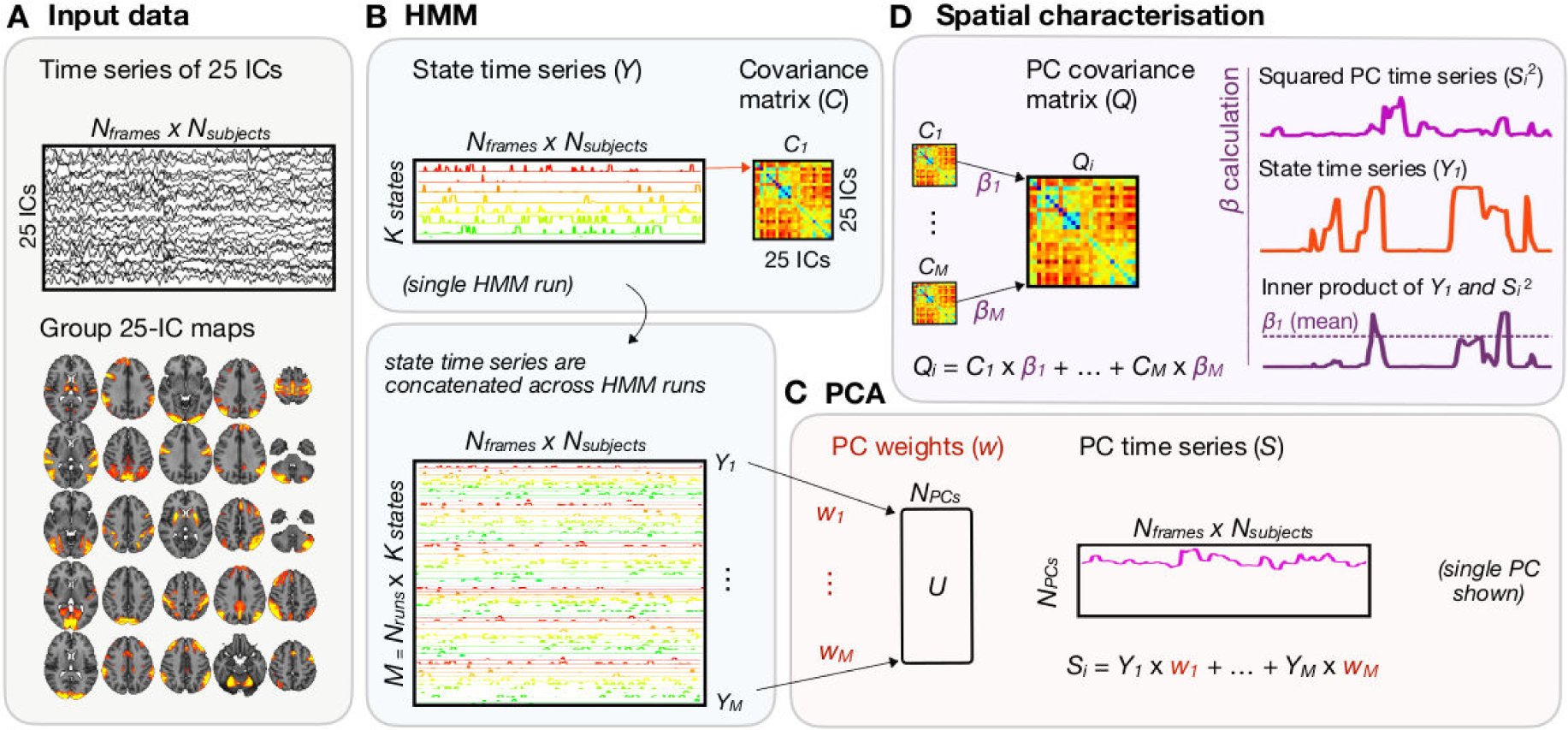
Graphical summary of PCCA. **A** processed BOLD timeseries of 25 ICs obtained from 100 subjects of the HCP; these timeseries were then standardised, separately for each subject. **B** HMM was applied to the concatenated BOLD timeseries across subjects (25 ICs by [*N_frames_* × *N_subjects_*]) resulting in *K* = 12 states, each with associated state timeseries (i.e., the probability of a given state to be active at each time point) and covariance matrix *C_m_* across 25 ICs. The HMM was run several times (N_runs_), each starting from a different random initialisation. **C** PCA was applied to the concatenated state timeseries across runs ([*N_runs_* × *K*] by [*N_frames_* × *N_subjects_*]), projecting the M (=*N_runs_* × *K*) rows into a reduced set of PCs explaining maximum variance. **D** The covariance matrix of each PC was estimated as a linear combination of the covariance matrices of the original (*M*) states, using the betas of each state (*β_m_*) as their weights. The betas were computed as the weighted average of each original state timeseries with the square timeseries of a given PC. HMM: hidden Markov model(ling); ICs: independent components; PCA: principal component analysis. PCCA: principal components of connectivity analysis.

To identify reliable patterns of time-varying FC, we developed PCCA, a novel approach designed to precisely characterise the time point by time point variability in FC. PCCA was run on each behavioural condition separately; see **Figure 1B-D** for a graphical representation of PCCA on the resting-state condition (and **Figure 1A** for a representation of the input data). In brief, PCCA consists of applying PCA (Jolliffe, 2002) to multiple runs of the HMM (Rabiner, 1989). Each HMM run included *K* = 12 states (**Figure 1B**); see *Reproducibility of the results* section for *K* = 6 and *K* = 16). These K states represent reoccurring patterns of FC. The probability of such FC pattern being active at each time point (i.e., state time series) is also estimated from the data. Since the HMM starts from a random seed and depends on an optimization procedure, different runs of the algorithm may result in somewhat different results. Here, we used this variability to our advantage by running the algorithm multiple times and performing PCA on the resulting state timeseries (**Figure 1C**). By doing so, we reduced the dimensionality of the data into a small set of orthogonal principal components (PCs) that capture the dominant fluctuations in FC, where each PC has an associated covariance matrix. The PC-specific covariance matrices were computed as weighted averages of the HMM state covariance matrices (**Figure 1D**); each weight, β, (one per HMM state and PC component) was calculated as the variance of a given PC timeseries during the time that the corresponding HMM state was active (see β calculation in **Figure 1D** and Methods for more details).

In what follows, we show that time-varying FC follows reliable trajectories that meet the three proposed criteria: that such trajectories are low but multi-dimensional, that their nature differs across behavioural conditions, and that FC exhibits changes within reasonably short timescales that are arguably compatible with the timescale of cognition and behaviour. In addition, we confirm that FC trajectories are not determined just by amplitude modulations. Finally, we verify the reproducibility of our results on new model configurations and a separate set of subjects.

### Functional connectivity reliably embeds into multidimensional temporal trajectories

We used our PCCA approach to capture modulations in brain network activity, in the sense of changing patterns of covariance across regions. By running the HMM multiple times, we leveraged the variability of the HMM inference by refactoring the different estimations into reliable, latent trajectories of time-varying FC. By doing so, we explored the complexity of these trajectories, finding that they robustly span multiple dimensions.

First, to quantify the stability of the separate instances of the HMM inference, we computed the Pearson correlation *r* between the state timeseries generated across multiple HMM runs. Because the ordering of the states within a run is arbitrary, the states were aligned between HMM runs using the Hungarian algorithm (Munkres, 1957), which, given two HMM runs, minimises a measure of cost given in this case by 1 minus *r*, summed across all pairs of matched states. By design, the states were ordered from best to worst aligned. We, therefore, obtained correlation coefficients *r* between every pair of aligned states. **Figure 2A** shows a histogram of the temporal *r* values (median = 0.71 ± 0.25) between all possible pairs of aligned states (19900 pairs). These results confirm that the states inferred by the HMM are to some extent different for the different runs of the inference (Vidaurre et al., 2019).

**Figure 2.**
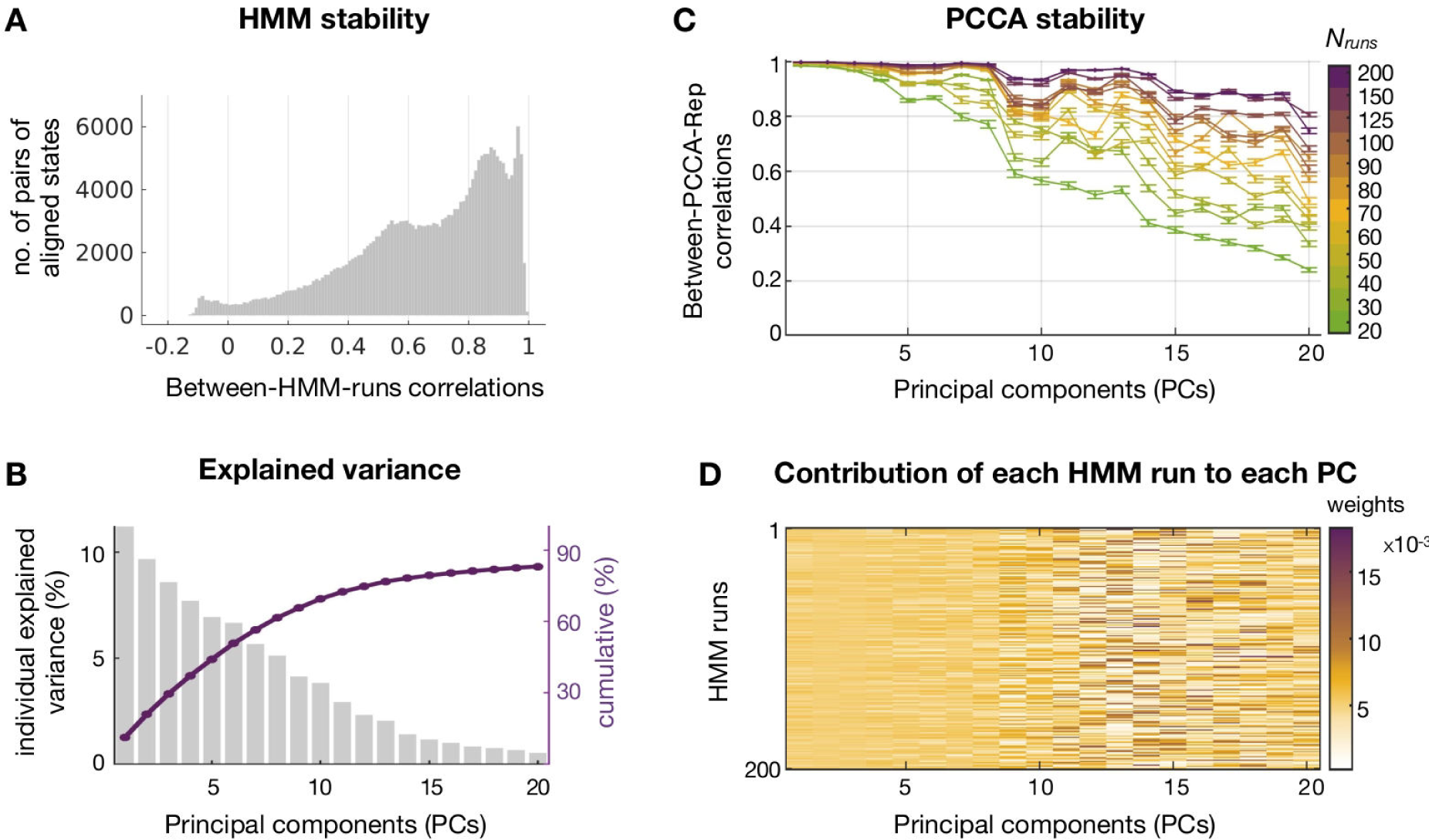
Resting-state time-varying FC spans multiple temporal dimensions that can be reliably measured. **A** Histogram of the Pearson correlation coefficients between (*K* = 12) aligned states for each pair of (200) HMM runs (19900 pairs in total). **B** Plot representing the percentage of individual (bars) and cumulative (line) explained variance by the top 20 PCs that resulted from applying PCA to the K = 12 states from 200 HMM runs (i.e., 12 × 200 = 2400 states). **C** Pearson correlation coefficient (y-axis) of the 20 PCs (x-axis) between each pair of 30 PCCA repetitions, for several numbers of HMM runs (from 20 to 200); error bars represent the standard error of the mean correlation across all pairs of PCCA repetitions. **D** Average state contribution (PCA weights) of each HMM run to the final PCCA model for the top 20 PCs.

As explained above and illustrated in **Figure 1B-C**, we exploited this variability by applying PCA across the state timeseries from 200 HMM runs. This produced a set of orthogonal PCs that optimally explained maximum time-varying FC variability across states and HMM runs (i.e., M=200 runs x 12 states per run = 2400 total states). As shown in **Figure 2B**, the top five PCs alone retained 44% of the total temporal variance present across all the states, suggesting that the temporal network variability in the data has more than five dimensions. Focusing on these five dimensions, we assessed the reliability of these PCs across 30 different PCCA repetitions, each of them consisting of *N_runs=_* 200 HMM runs with K=12 states per run. We then estimated the reliability of each PC by computing the Pearson correlation *r* between the PC timeseries of each pair of PCCA repetitions. The average across all pairs was *r* ≈ 1.0 for PC1 and PC2, and *r* ≈ 0.99 for PC3, PC4 and PC5, confirming that our approach is extremely stable against estimation noise —of which other approaches suffer systematically (see Hindriks et al., 2016; Vidaurre et al., 2019). To further quantify the relation between the number of HMM runs (*N_runs_*) and the stability of the results, we repeated this process for various choices of *N_runs_*. **Figure 2C** shows that the set of states generated over 30 PCCA repetitions was already able to make the five PCs highly stable (*r* > 0.92). While this stability decreased for lower-order PCs, adding more HMM runs did increase the number of stable PCs. Additionally, since the aim of applying PCA was to integrate information across HMM runs, we verified that the PCs were not representations of specific runs. As observed in **Figure 2D** the average state contribution of each HMM run to the PCs are similar across runs (particularly concerning their contributions to higher-order PCs), demonstrating that the PCs effectively represent a low-dimensional embedding across all runs.

Similar conclusions were drawn from the analysis of task fMRI data, as depicted in **Figure S1**: during the WM and the motor task conditions, FC followed reliable trajectories across multiple dimensions. Compared to the resting-state condition, there is a higher variability between the states of different HMM runs during the task conditions, particularly during the WM task. Due to this increased variability, more HMM runs were thus needed to ensure stable PCCA results for the task conditions. However, similar to the resting-state condition, the top five PCs were highly stable during both task conditions (*r* > 0.9); these retained around 40% of the total variance present across the 2400 states, suggesting that FC organises into even higher dimensions.

The implications of these results are two-fold. First, despite the stochastic nature of the HMM, momentary changes in FC can be reliably captured. Second, the fact that not only one but several FC dimensions were required to explain half of the total variance in the data, and that the first dimension alone only explained around 11% of the total variance, demonstrates that fluctuations in FC are low but multi-dimensional. For convenience, we hereon focus on these top five dimensions (unless explicitly stated).

### FC fluctuations depend on the behavioural context

After extracting reliable fluctuations of FC using PCCA, we can now investigate how the spatial pattern associated with these fluctuations vary across behavioural context (i.e., rest, WM, and motor). If time-varying FC reflects the dynamic configurations by which brain regions interact to shape cognition, we expect the spatial maps of FC fluctuations to be context-dependent. Here we examined the spatial patterns of the estimated FC trajectories for the top 20 PCs (instead of five, for completeness). Specifically, we transformed the covariance matrices of the 20 PCs of each behavioural condition into correlation matrices and then applied the Fisher z-transformation on the off-diagonal elements of the matrices; these were then vectorised and compared to one another using Pearson correlation. All correlation coefficients are provided in **Figure 3A**. To determine whether the overall correlation within and between conditions was significantly different, we used permutation testing (10000 permutations). The significance level was adjusted using false-discovery rate (FDR; Benjamini and Hochberg, 1995). **Figure 3B** provides an overview of the correlation differences. See brain connectivity graphs and connectivity maps of the top five PCs for each behavioural condition in **Figure S2**.

**Figure 3.**
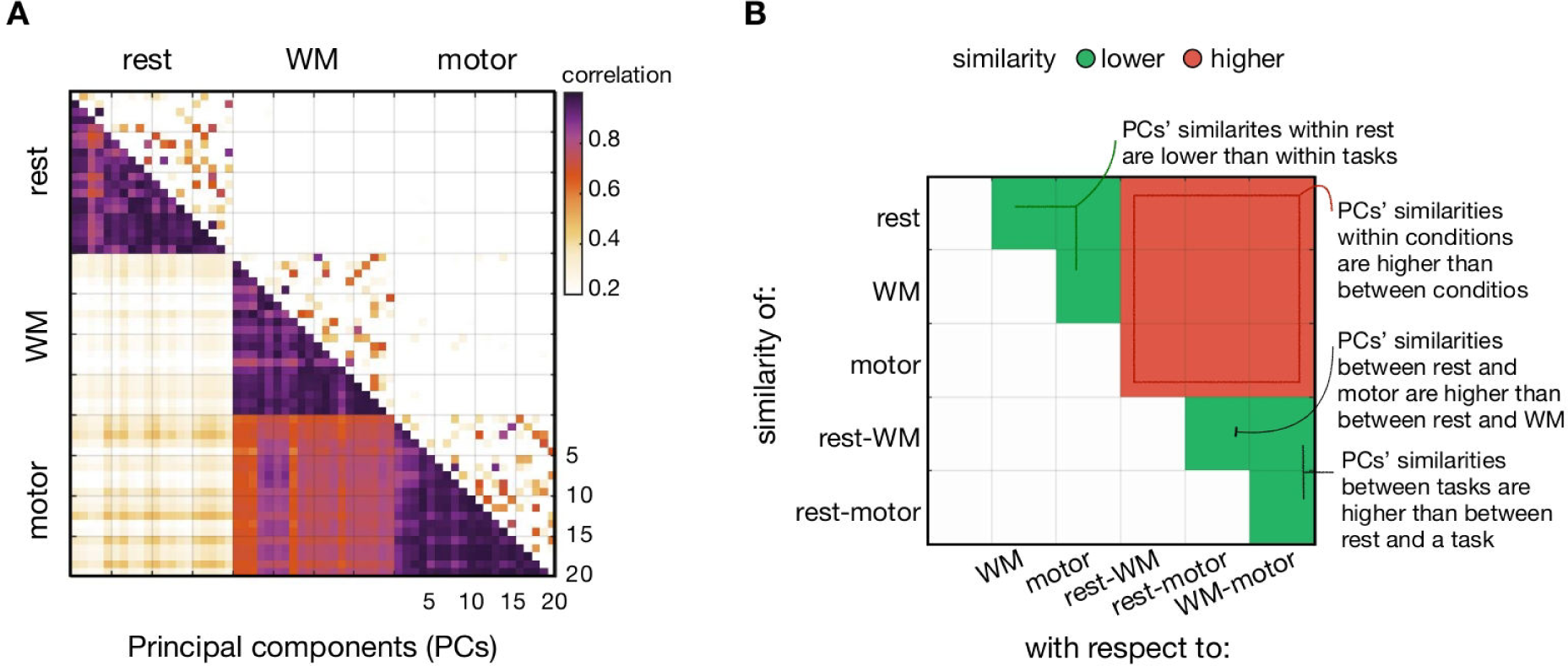
The spatial patterns of the FC trajectories are condition specific. **A** Correlation (lower triangular) and partial correlation (upper triangular) coefficients across the top 20 PCs within and between conditions: rest, WM, and motor (each row and column are a different PC). **B** Significant (p-FDR < 0.05) differences (lower in green and higher in red) of the averaged similarity of the spatial patterns (of rows with respect to columns) for all pairs of within (rest, WM, motor) and between (rest-WM, rest-motor, WM-motor) conditions. Statistical analysis based on permutation testing (10000 permutations).

First, we found that changes in FC were condition-specific, that is, they depend on the behavioural context. As indicated by the red block in **Figure 3B**, the overall spatial similarity across PCs (i.e., the averaged correlation across the top 20 PCs) was significantly higher within than between conditions (*p-FDR* < 0.0001). Second, the largest differences were observed between the rest and task conditions, i.e., the similarity of the spatial patterns of FC fluctuations between the two tasks was always higher than between a task and the rest condition (*p-FDR* < 0.0001; lower green triangle in **Figure 3B**). Although the WM and motor tasks were designed to target different cognitive processes (i.e., working memory function and motor execution, respectively), there are common aspects to the two tasks such as motor coordination and visual processing, and potentially other factors related to attention (e.g., following instructions), that might be causing the resting state to be more markedly different. Third, we observed that the differences between rest and task PCs were even more pronounced for the WM than for the motor task (*p-FDR* < 0.0001), which could partially be explained by the higher experimental demands of the WM task compared to the motor task. Fourth, we found that the spatial patterns of FC fluctuations were far more diverse (i.e., lower correlation across PCs) during rest than during task conditions (*p-FDR* < 0.01; upper green triangle in **Figure 3B**), probably due to the higher capacity of the participants to engage in a more unconstrained type of cognition during the resting-state (Wang et al., 2018).

In summary, these results indicate that the dominant modulations in FC meaningfully relate to the behavioural context and that these modulations are more diverse during rest than during task conditions —*possibly* reflecting the unconstrained nature of the resting state.

### FC fluctuates at timescales that are compatible with cognition

Having shown that the nature of the estimated FC trajectories is context-dependent, we sought to investigate the temporal scales of these modulations. We reasoned that for FC to relate to dynamic neural communication, FC should exhibit changes at timescales that are compatible with the timescales of ongoing cognition. Focusing on the memory task, we asked how differences in WM variables (i.e., WM performance and WM load), which vary across blocks, would relate to each of the different timescales underlying the FC trajectories.

To dissect the various FC-related timescales, we used the empirical mode decomposition (EMD, Fabus et al., 2021; Huang et al., 1998), a data-driven approach that allowed us to split each FC trajectory (i.e. each PC timeseries) into different so-called intrinsic mode functions (IMFs). Each IMF covers a certain range of frequency, with one instantaneous frequency value per time point. Focusing on the top five PCs, the EMD yielded ten IMFs for each PC timeseries (see **Figure 4A** and **Figure S3**), where the first IMF (IMF1) corresponded to the fastest scale (with the range 0.03Hz to 0.31Hz covering 90% of the instantaneous frequency values) and the last IMF (IMF10) to the slowest scale (90% range of 0.00003Hz to 0.0001Hz).

**Figure 4.**
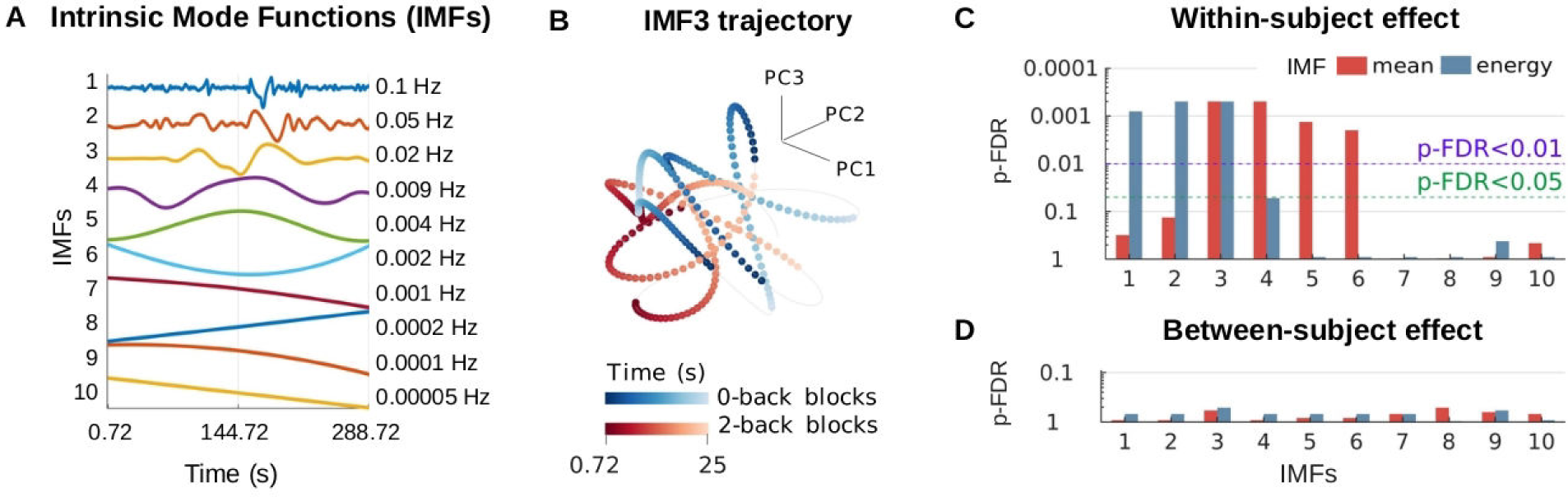
Fluctuations in FC are associated with cognitive features. **A** Illustration of IMFs for a section of the data; the labels on the right indicate the median instantaneous frequency of each IMF averaged across subjects and (top five) PCs. **B** projection of IMF3 trajectory for a single subject’s session, with colours representing the periods of 0-back (in blue) and of 2-back (in red) blocks; the intensity of the colours indicates the beginning and the end of each block (25s). **C** Bar plot shows the FDR-corrected p-values of the canonical correlation between cognitive abilities (i.e., WM load, RT and accuracy) and two descriptive statistics of IMF-specific PC time series —i.e., IMF mean (red) and IMF energy (blue). **D** Bar plot shows the FDR-corrected p-values of the canonical correlation between cognitive abilities (i.e., RT and accuracy) within 2-back blocks and two descriptive statistics of IMF-specific PC time series —i.e., IMF mean (red) and IMF energy (blue); p-values were obtained using permutation testing: permutations were performed within subjects in C, and between subjects in D. All p-values were FDR corrected across IMFs and descriptive statistics (i.e., 10 x 2 tests in C and 10 x 2 tests in D).

To assess whether the dominant fluctuations in FC within a given frequency band (i.e., IMF-specific PC timeseries) relate to cognition (expressed here as block-specific WM variables), we used canonical correlation analysis (CCA; Wang et al., 2020) and permutation testing (10000 permutations) using the canonical correlation coefficient as the base statistic. The association between WM variables and IMF-specific PC timeseries were run separately for each IMF. The set of WM variables was encoded as a three-column matrix, containing the average accuracy, the average reaction time of correct trials, and the WM load of each block. These were concatenated across sessions and subjects (i.e., 8 blocks x 2 sessions x 99 subjects = 1584 rows). WM load was a zero-one indicator variable with the zero indicating 0-back blocks and one indicating 2-back blocks. The set of IMF-specific variables was encoded as a five-column matrix containing, for the top five PCs, a descriptive statistic (either the mean or the variance) of the period of the PC timeseries corresponding to each of the 1584 blocks. While the mean IMF-specific PC timeseries within a block captures differences in phase of these EMD-derived FC trajectories, the variance of the IMF-specific PC timeseries can be conceptualised as the “*energy*” of these FC trajectories. Separate tests were run to test each descriptive statistic. FDR was used to correct across multiple comparisons (2 descriptive statistics x 10 IMFs = 20 tests). Critically, in order to focus on within-session differences, permutations were performed within each subject across blocks and sessions but not across subjects. See **Table 1** for details of the WM task design. For illustration, **Figure 4B** projects the IMF3 trajectories of the top three PCs for a single subject’s session, showing periods of 0-back blocks (in blue) and 2-back blocks (in red). IMF3 (median frequency = 0.02Hz; 90% range of 0.01Hz to 0.04Hz) was chosen since both its mean and energy exhibited a significant (p-FDR < 0.001) relation to WM. The mean of IMF1 and IMF2, and the energy of IMF4, IMF5 and IMF6, were also significant. As we expected, we see phase effects in lower frequencies and energy effects in larger frequencies (**Figure 4C**).

**Table 1.** Data description

| Condition | Subjects | Sessions | Frames<br>per<br>session | Description |
| --- | --- | --- | --- | --- |
| Rest | 100 | 4 | 1200 | Participants were instructed to try to lie still, to keep their eyes open (relaxed fixation), not to think of anything specific, and not to fall asleep (although some fall asleep at some point). |
| Working<br>Memory<br>(WM) | 99 | 2 | 405 | Participants were presented with pictures (places, faces, body parts and tools) in alternating blocks of 0-back and 2-back WM. During 0-back blocks, participants had to respond to any presentation of a target cue (presented at the start of the block). During 2-back blocks, participants had to respond to whether the current stimulus was the same as the two stimuli earlier. |
| Motor | 100 | 2 | 284 | Participants were presented with visual cues that required them to tap their left or right fingers, squeeze their left or right toes, or move their tongue. |

In a secondary analysis, we looked at between-subject effects by assessing the canonical correlation between IMF-specific PC time series and WM variables (i.e., accuracy and reaction time) for the 2-back blocks. Accounting for the family structure of the HCP data, permutations were performed between subjects instead of between blocks(Smith et al., 2015; Winkler et al., 2015). As observed in **Figure 4D**, we did not find statistical evidence for a relationship between the subject’s performance on the 2-back blocks and the mean or energy of the PC timeseries within any of the EMD-estimated frequency bands. The absence of statistical significance when assessing between-subject effects may be related to the low sample size (Marek et al., 2022, see *Discussion*). **Table S1** and **Table S2** show the CCA results in more detail for the first (i.e., within-session) and second (i.e., between-subject) analysis respectively.

Overall, these results suggest that FC modulations can capture distinct cognitive processes imposed by the task (i.e., low vs high WM load), revealing that various timescales of the estimated FC fluctuations (IMF1– IMF6) relate to behavioural information such as cognitive abilities and cognitive demands. Although there are much faster dynamics that cannot be captured by fMRI, the estimated FC modulations fluctuate within a session at velocities (i.e., from seconds to tens of seconds) that are compatible with at least some aspects of human cognitive processing.

### Functional connectivity trajectories are not purely driven by simple activation patterns

Since we are interested in whether dynamic cross-regional communication can be related to FC trajectories, we sought to verify that such trajectories are not determined by just amplitude modulations. To disambiguate whether the PC modulations contain information that is unique to time-varying FC (i.e., beyond just regional activations and amplitude modulations), we examined how much of the temporal variation in FC can be predicted by changes in the raw BOLD signal (represented as 25 IC timeseries). Using cross-validated regularized ridge regression (Hoerl and Kennard, 1970) on each PC separately, we show that the main patterns of temporal fluctuations in FC during rest cannot be well explained by changes in order-1 activations (**Figure 5**). Specifically, the IC timeseries can only explain 1% of the variance of the top four PCs and 11% of the fifth PC. In contrast, the order-2 interactions of the signal (i.e., the pairwise product of the IC timeseries, which is directly related to Pearson-based FC), explained considerably more variance of the PCs (values ranging between 18% and 43%). Note that because the predictions are cross-validated, this comparison is not biased by the fact that the number of order-2 interactions (300 timeseries, one per pair of ICs) is larger than the number of order-1 activations (25 IC timeseries). Moreover, since the order-2 interactions also contain the information of the amplitude itself (within the diagonal of the covariance matrices), we repeated the prediction after regressing out the raw signal of the PC timeseries in a cross-validated fashion (Snoek et al., 2019). On average, less than 2% of the explained variance was lost after accounting for the raw signal, indicating that the PCs do contain information that is unique to time-varying FC.

**Figure 5.**
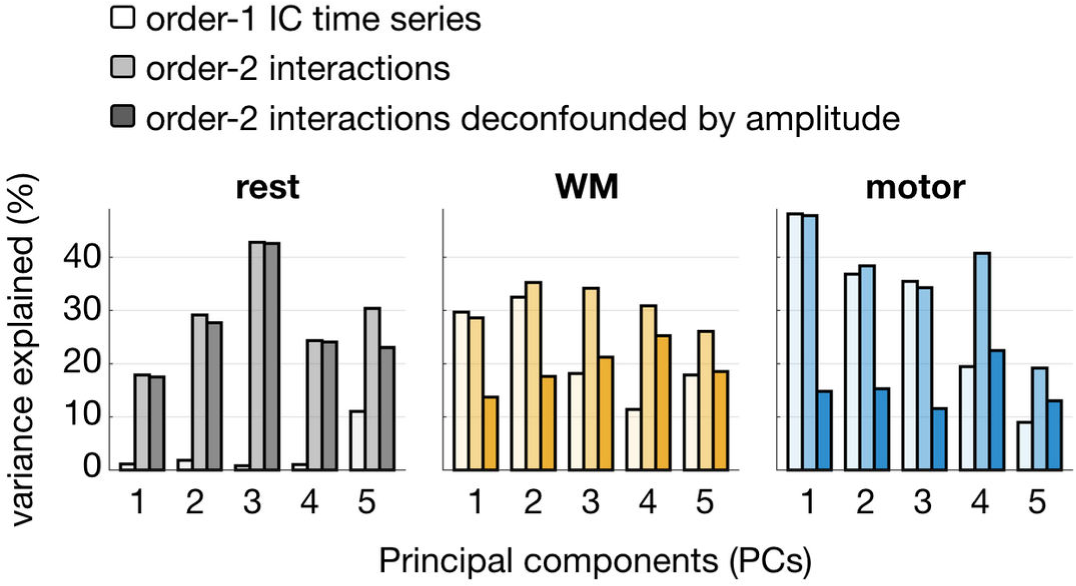
PCCA timeseries cannot be fully explained by amplitude. Percentage of explained variance (R2, y-axis) of the top 5 PCs (x-axis) by the (order-1) IC timeseries, and order-2 interactions with and without deconfounding for amplitude. Components are coloured according to 3 conditions: rest, WM, and motor.

During the task conditions, however, the changes in order-1 timeseries contributed more to explaining the fluctuations in FC. This was expected since the evoked responses are synchronised across brain regions by the task experimental design, creating task-induced correlations and therefore contributing to the FC estimations. This effect was more pronounced for the motor task than for the WM task condition. Specifically, the IC timeseries could explain up to 48% of the variance of the first PC during the motor task, and 32% of the second PC during the WM task. Order-2 interactions of the signal had similar contributions to the amplitude for the first two PCs of the WM task and the first three PCs of the motor task. For the remaining PCs, order-2 interactions explained more variance than the IC timeseries. As before, we repeated the prediction after correcting for the order-1 activations, finding that the variance explained by order-2 interactions decreased around 12% for the WM task, and 21% for the motor task, after deconfounding for the IC timeseries. This loss of variance indicates that a considerable proportion of the fluctuations in FC are driven by changes in signal amplitude that are likely evoked by task responses.

In conclusion, these results demonstrate that there exists unique information contained in the PCCA trajectories that cannot be fully predicted by regional activation.

### Reproducibility of the results

We sought to evaluate the reproducibility of our results with respect to five different variations of the experimental setup (**Figure 6**, left to right): using a separate set of subjects, using the original dataset with different model configurations (changing the fixed number of HMM states from 12 to 6 and to 16), and using different data configurations (reducing the length of the timeseries from 4800 to 810 frames; using a subsample with half of the subjects). We focused on the resting state, free of task-related activations that could trivially explain the reproducibility scores and used the top 20 PCs (instead of five) for completeness.

**Figure 6.**
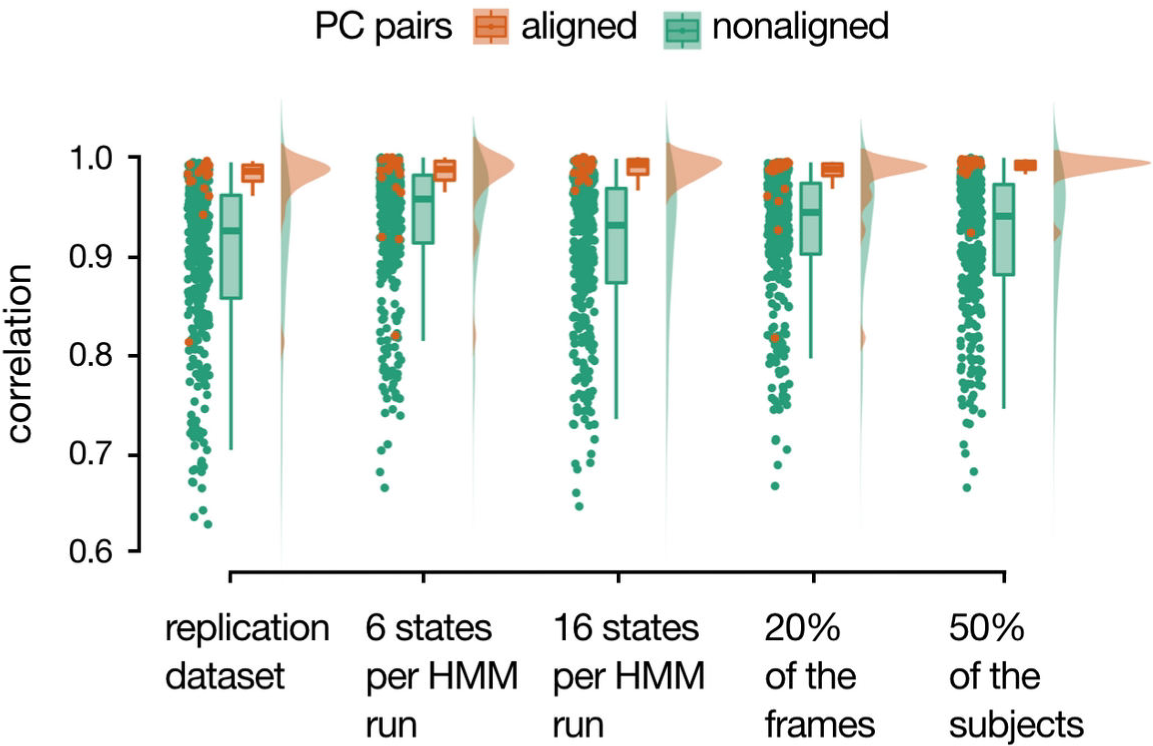
Reproducibility of the time-varying FC patterns for the top 20 PCs in the resting-state. From left to right, reproducibility of our results on a separate set of subjects (i.e., 100 subjects, 4600 frames, 12 states), and with respect to changes in the number of states (from 12 to 6 and 16 per HMM run), to different scan lengths (from 4600 frames to 810), and different sample sizes (from 100 to 50 subjects). Box plots represent the inter-quartile range (IQR; between the 75^th^ and 25^th^ percentiles) and median (horizontal line) of the correlation values (one per dot) between aligned PCs (orange) and non-aligned PCs (green).

For each of these five experimental variations, we first extracted the top 20 PCs following the same procedure as described in **Figure 1**. As discussed, we transformed the covariance matrices of each of these 20 PCs into correlation matrices, applied the Fisher z-transformation on the off-diagonal elements of the matrices and vectorised them. These vectors were then compared to those of the original dataset using Pearson correlation. Reproducibility was assessed by comparing whether the correlation between aligned PCs was significantly higher than between non-aligned PCs. The statistical significance of the difference was determined using permutation testing (significance level of 0.05; 10000 permutations).

We found that the overall changes in FC (as captured by the top 20 PCs) were highly reproducible for all five experimental variations. That is, the correlation between aligned PCs was significantly higher than between non-aligned PCs (*p* < 10^-5^). When using a separate set of subjects (first column of **Figure 6**) only three out of 20 aligned PCs (orange) fall below the 75^th^ percentile of the correlations between non-aligned PCs (green). Concerning changes in model configurations, reducing the number of fixed states from 12 to 6 (second column of **Figure 6**) returned 14 very similar PCs; while increasing it from 12 to 16 (third column of **Figure 6**) returned 19 very similar PCs, suggesting that our approach is robust to the number of HMM states within a reasonable range. When reducing the length of the timeseries to 810 frames (the length of the WM task), 15 of the 20 PCs could still be recovered even though only a fifth of the original number of volumes were available (fourth column of **Figure 6**). Lastly, only one of the 20 PCs could not be recovered when estimating the PCCA model on just half of the subjects from the original dataset (fifth column of **Figure 6**).

## Discussion

Can the temporal variability of patterns of fMRI FC across multiple regions reflect information transfer between the underlying neuronal populations, or do they rather *just* reflect stable anatomical pathways? We propose that, for the former case to be plausible, these correlations should fulfil, at least, the following criteria: (i) its latent complexity must be relatively high to accommodate the flexibility that brain communication requires; (ii) it should modulate its configurations according to the behavioural context; and (iii) it should exhibit changes at timescales that are compatible with the timing of cognition. Note that, while for example structural connectivity also in some sense reflects neural communication, we here refer to actual neural communication mechanisms subserving cognition, which are necessarily dynamic. Using PCCA, a novel approach based on multiple runs of the HMM in combination with PCA, we showed that reliable estimations of FC patterns can be represented as low-dimensional trajectories of order-2 interactions. Importantly, these could not be fully explained from order-1 signal activations. These FC-related trajectories were found to meet the proposed criteria, suggesting that they may faithfully reflect at least some aspects of large-scale, cross-regional communication.

In this paper, we have analysed conditions that are required for FC to reflect neuronal communication. With present data, however, it cannot be proven categorically that these conditions are sufficient, and that time-varying FC univocally reflects large-scale communication and, thus, cognition. Indeed, the dynamics of BOLD signals and their covariance are, at least partially, modulated by non-neural factors such as respiratory cycles, cardiac activity, arterial CO_2_ concentration, and, of course, motion (Murphy et al., 2013; Power et al., 2017; Xifra-Porxas et al., 2021). Importantly, while these are not neural effects per se, most of these factors are inextricably linked to actual neural activity. Our view is that, while the estimated PCCA trajectories are not independent of these non-neural factors, they are likely to also reflect neural communication dynamics. However, since these distinct factors —neural and non-neural— are coupled in highly non-linear ways, caution must be exercised in their interpretation. That is, saying that FC can index neural communication does not mean that this is a direct relationship, or that this modulation does not occur through several other factors.

The fact that not only one but several FC dimensions could be reliably estimated might seem at odds with previous work where network activity was shown to be hierarchically organized into two groups of “metastates” (Vidaurre et al., 2017). Indeed, having the states strongly clustered into two “metastates” essentially defines a one-dimensional axis of variation for FC. This discrepancy can be explained twofold. First, in Vidaurre et al. (2017) the HMM was run only once, whereas here we have used multiple runs; this allowed us to explore the space of solutions exhaustively. Second, and most importantly, while in the previous study PCA was applied on the fractional occupancies (an aggregated measure where each session is defined as how much time the subject spent on each state), here we have applied PCA to the state timeseries. That is, in the previous study PCA was performed on a (*no. of subjects* by *no. of states*) matrix, and here PCA is applied on a much larger ([*no. of subjects* × *no. of time points* × *no. of sessions*] by [*no. of states* × *no. of HMM runs*]) matrix. Therefore, whereas the “metastates” in Vidaurre et al. (2017) fundamentally reflect between-subject variability of *both* time-varying and time-averaged FC, PCCA characterises the actual temporal dimension of the FC patterns, where multiple dimensions are shown to be at play.

To understand the nature of FC modulations and their relationships to cognition and behaviour, we have investigated to what extent the estimated FC-related trajectories reflect changes in the environment by looking at different tasks. We examined the spatial correspondence of the estimated FC trajectories across different behavioural contexts: at rest, during a WM task, and a motor task. On the one hand, our analysis revealed distinct spatial patterns for each of the three conditions, indicating the behavioural specificity of the estimated trajectories. These differences are particularly important since they emphasise the dynamic configurations in which brain regions’ interactions evolve. On the other hand, we found a higher similarity of brain network modulations during the execution of tasks compared to rest, highlighting the constrained element of both WM and motor tasks as opposed to the unconstrained nature of the resting-state. A possible explanation is that the task designs elicit patterns of synchronised activity that are to some extent generalisable across tasks (for instance related to attentional processes), and these become reflected in the network trajectories. This observation aligns with previous evidence that task-induced changes in FC are not simply driven by task-evoked activation (Greene et al., 2020). Recent studies analysing how brain activity unfolds within a low-dimensional space have also found common patterns across several tasks (Saggar et al., 2018; Shine et al., 2019a), and observed that modulations in these patterns have been associated with differences in task demands and task performance (Cornblath et al., 2020; Saggar et al., 2018; Shine et al., 2019b). As opposed to our PCCA approach, these latent trajectories were computed based on first-order statistics and are therefore unspecific to connectivity. Our work extends these findings by demonstrating that within-session variability in cognitive demands and cognitive abilities shape the temporal evolution of brain FC (and not just activity) at various timescales. Moving forward, investigating the different timescales at which FC fluctuate may offer new informative paths in the future about its role in cognition. Noteworthy, we did not find a reliable mapping between inter-individual differences in WM performance and FC fluctuations, which could be attributed to having a too low sample size (Marek et al., 2022).

These results emphasise the importance of neural variability for behaviour (Waschke et al., 2021), and align with previous reports on the variability in BOLD signals as an important feature of brain function (Garrett et al., 2020, 2010; McIntosh et al., 2008). Using EEG and fMRI, these studies showed that higher brain signal variability facilitates the formation of brain networks and transitions between them, which in turn enhances the brain’s dynamical repertoire to enable cognitive function. Complementing these studies, we quantified the degree of variability in the space of brain network interactions, therefore extending the notion of signal variability towards connectivity variability. Furthermore, we found brain network modulations to be more diverse during rest than during tasks, which has been consistently reported in the literature (Chen et al., 2015; Elton and Gao, 2015; Hutchison and Morton, 2015). This finding appears to reflect the unconstrained nature of the resting-state and supports the idea that resting-state FC may provide a richer characterization of brain activity than task states (Ponce-Alvarez et al., 2015).

Previous research on time-varying FC has often identified a set of brain states occurring within the full scanning session (Lurie et al., 2020), for example using an HMM (Vidaurre et al., 2018) or by performing (e.g., k-means) clustering on sliding-window estimates (Allen et al., 2014). Despite the specifics of each approach, the choice of the number of states is a cross-methodological concern. Here, we found that irrespective of the initial number of fixed states (*K* = 6, 12 and 16), PCCA was able to reliably capture the same prevalent underlying configurations. The highly robust estimation across choices of *K* indicates that the relative instability of the HMM inference can be turned into an advantage when harnessed by PCA; see Vidaurre et al. (2019) for a similar argument in the context of statistical testing. Methodologically, PCA might offer a promising alternative to conventional model selection, toward model integration.

A limitation of the present method is that, if we run the HMM multiple times and we find variability between the models, this variability can be due to two causes: the complexity of the landscape of FC dynamics, which will induce estimation variability since the one HMM’s explanatory power is not sufficient; and pure estimation noise, due to factors such as not enough data or high signal-to-noise ratio. While this is not a problem regarding the interpretation of the results, it can be if we wish to compare the complexity of the landscape of FC dynamics between two conditions (e.g., rest and a motor task) with different levels of noise and amount of data. Although not explored in this paper, these two sources of variability might potentially be disentangled by looking at the PCA eigenspectra patterns. This will be explored in future work. Another avenue for improvement is to explore alternatives for PCA that might be better suited for time series of probabilities (Lee et al., 2010). Also, note that both the HMM and PCA are run at the group level, meaning that while the HMM and PCCA time series are estimated at the subject level, all the spatial estimates are at the group level. Future approaches could aim at obtaining a hierarchical estimation to boost between-subject or between-condition differences (similarly to Harrison et al. 2015). Finally, note that, while we have used a resting-state (data-driven) parcellation and then applied it on task, it would have also been possible to derive the parcellation from task data and then apply it to the resting-state data. We opted for the more standard approach of using the resting-state as an initial point because, as shown above, the resting-state dynamics are more diverse than the task-state dynamics and reflect the landscape of FC dynamics more broadly.

In summary, our results indicate that brain network modulations fluctuate at relatively short timescales following reliable patterns that are context-dependent, influenced by cognitive complexity and associated with cognitive abilities. The estimated trajectories are relatively low-but still multi-dimensional, meeting the minimum requisites to reflect the dynamics of information transfer across the brain.

## Methods

### Data Description

We used publicly available fMRI data from 100 subjects from the HCP (Van Essen et al., 2013). For each subject, we considered data from the resting-state (rest), the WM task, and the motor task. One subject’s WM task data was missing. Brief descriptions of each condition are provided in **Table 1**. Please refer to Van Essen et al., 2012 for full details about the acquisition and preprocessing of the data. In brief, 3T whole-brain fMRI data was acquired with a spatial resolution of 2×2×2 mm and a temporal resolution of 0.72 s. All fMRI processing was performed using FSL (Jenkinson et al., 2012) including minimal highpass temporal filtering (>2000s FWHM) to remove the linear trends of the data, and artefact removal using independent component analysis (ICA)+FIX (Griffanti et al., 2014; Salimi-Khorshidi et al., 2014). No lowpass temporal filtering or global signal regression was applied. Group spatial-ICA was performed using MELODIC (Beckmann and Smith, 2004) on the resting-state data in order to obtain a parcellation of 25 ICs. Whereas a higher number of IC parcellations are provided by the HCP, the 25-ICA parcellation was sufficient to map the dynamics of FC as it covers the major functional networks, while keeping the computational cost relatively low. For ease of comparison across conditions, the timeseries for each IC were computed using the resting-state group 25-ICA parcellation for each subject and condition (rest, WM, and motor) through dual regression (Filippini et al., 2009). Specifically, the IC timeseries corresponding to the task sessions were computed from the fully preprocessed data in MNI152 space as delivered in the HCP1200 release; the IC timeseries of the resting-state sessions were directly obtained from the HCP ‘PTN’ Parcellation+Timeseries+Netmats (specifically, the first 100 subjects from the ‘recon2’ version).

### Principal Component of Connectivity Analysis

PCCA is composed of two methodological elements: the HMM and PCA. The parameters of PCCA to be chosen by the user are the number of HMM states and the number of HMM runs. We next describe these, including how to compute connectivity maps from the resulting principal connectivity components.

### Hidden Markov modelling

The HMM was used to extract whole-brain patterns of time-varying FC during the rest and task conditions. In a data-driven way, the HMM characterises neural timeseries of concatenated data using a finite number of *K* (group-level) states that reoccur across time (Rabiner, 1989), where the *k*-th state timeseries represents the probability for that state to be active at each time point; that is, the spatial parameters of the HMM are at the group level, while the temporal parameters are at the subject level. The states themselves are probability distributions within a certain family, and each is characterised by a certain set of parameters. To focus on FC, each state was here characterized by a Gaussian distribution with no mean parameter (the mean parameter pinned to zero) and a full covariance matrix (representing pairwise covariations across regions); or, equivalently, as a Wishart distribution (Vidaurre et al., 2021). Finally, a *K-*by-*K* transition probability matrix is estimated as part of the model, containing the probability of transitioning from one state to another or remaining in the current state. Here, the number of *K* states was set to 12. Note that since there is no specific biological significance in the number of states, this number was chosen to be consistent with our previous applications of HMM to fMRI data (Vidaurre et al., 2017, 2019). The HMM was applied to the concatenated (standardised) timeseries for all subjects ([*N_runs_* × *K]* by [*N_frames_* × *N_subjects_*]) for each condition separately (rest, WM task and motor task; see **Table 1**). See **Figure 1A-B** for a schematic description of the HMM.

### Principal component analysis and component characterization

PCA was applied to the HMM state timeseries to integrate the time-varying FC estimations across many runs. In short, the state timeseries generated over multiple runs (*N_runs_*) were attached to a matrix (*Y*) with as many rows as frames in the data set (*N_frames_* × *N_subjects_*) and *M* columns (*N_runs_* × *K* states); see **Figure 1B-C**. PCA was then performed by eigendecomposing *Y Y*^T^, which returns a set of *M* eigenvectors and eigenvalues according to the equation:

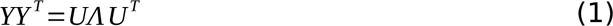

where *U* contained the (orthonormal) eigenvectors on its columns and Λ contained the corresponding eigenvalues in its diagonal (**Figure 1C**). These eigenvectors, or PCs, represented patterns of variation across all time-varying FC states, while the eigenvalues described the contribution of each PC in terms of explained variance. The PCs (columns of *U*) were arranged according to their relative contributions to *Y*, from higher to lower eigenvalues.

To estimate the spatial patterns of time-varying FC associated with the *i*-th PC, we computed one covariance matrix for each PC (denoted as *Q_i_*) by weighting the covariance matrices of the original states (*C_m_*) with weights β*_m_* as follows:

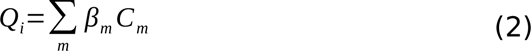

The weights β*_m_* were calculated as the variance of the (1 by [*N_frames_* × *N_subjects_*]) PC timeseries *S_i_* when state *m* is active:

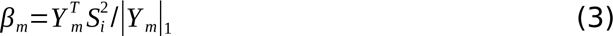

where *S_i_* represents the timeseries of the *i*-th temporal PC (estimated at the subject level), *Y_m_* is the probability of that state to be active or not at each time point and subject (**Figure 1D**), and |*Y_m_*|_1_ represents the sum of all elements of *Y_m_* (i.e., the L^1^ norm of the vector). Note that it is not possible to estimate the PCs’ covariance matrices using the PCA weights directly, because these are not necessarily positive and therefore the resulting matrix would not necessarily be positive definite (a mathematical requirement for any covariance matrix). Therefore, Eq. (3) is defined by convention and does not follow first principles.

To facilitate visualization of the spatial pattern associated with each PC, we used the first eigenvector of the corresponding PC covariance matrix (*Q_i_)*, which represented the main pattern of FC for that PC. Note that this is a different eigendecomposition than Eq. (1), which is done on the PC timeseries. The brain connectivity maps shown in **Figure S2** were created by overlaying these eigenvectors on brain surface space.

### Empirical mode decomposition

Using an iterative algorithm, the EMD aims at decomposing multi-frequency timeseries into a number of oscillatory modes referred to as IMFs, where each IMF relates to one fundamental frequency (Huang et al., 1998; Quinn et al., 2021). Here, we used the (univariate) EMD to decompose each FC trajectory (i.e., PC timeseries) into a set of IMF-specific timeseries of the same length as the PC timeseries. This was done for each PC timeseries separately. Note that the EMD was run on the state timeseries from the HMM estimation (which is a non-linear transformation of the data) and not on the BOLD timeseries. As a result, we might find IMF fluctuations slower than the filtered BOLD timeseries. The number of IMFs was determined in a data-driven way by the intrinsic temporal-spectral characteristics of the PC timeseries. The decomposition of the PC timeseries yielded ten IMFs (IMF1–IMF10), where each IMF covered a certain frequency range; see **Figure S3** for the range of frequencies included in each IMF and the relative energy across frequencies within and IMF. The instantaneous frequencies and energies of each IMF were obtained using the Hilbert-Huang Transform (Huang et al., 2009).

## Code availability

The code used to analyse the data in this study comprises our open-access code repository (https://github.com/OHBA-analysis/HMM-MAR) and custom scripts that are publicly available at https://github.com/sonsolesalonsomartinez/PCCA.

## Acknowledgements

SA and GH were funded by a donation of Hazewinkel. AL has been supported by the Horizon2020 programme CANDY Grant (No. 847818). DV is supported by a Novo Nordisk Foundation Emerging Investigator Fellowship (NNF19OC-0054895) and an ERC Starting Grant (ERC-StG-2019–850404).

Data were provided by the Human Connectome Project, WU-Minn Consortium (Principal Investigators: David Van Essen and Kamil Ugurbil; 1U54MH091657) funded by the 16 NIH Institutes and Centers that support the NIH Blueprint for Neuroscience Research; and by the McDonnell Center for Systems Neuroscience at Washington University.

## Supporting Information

**Figure S1.**
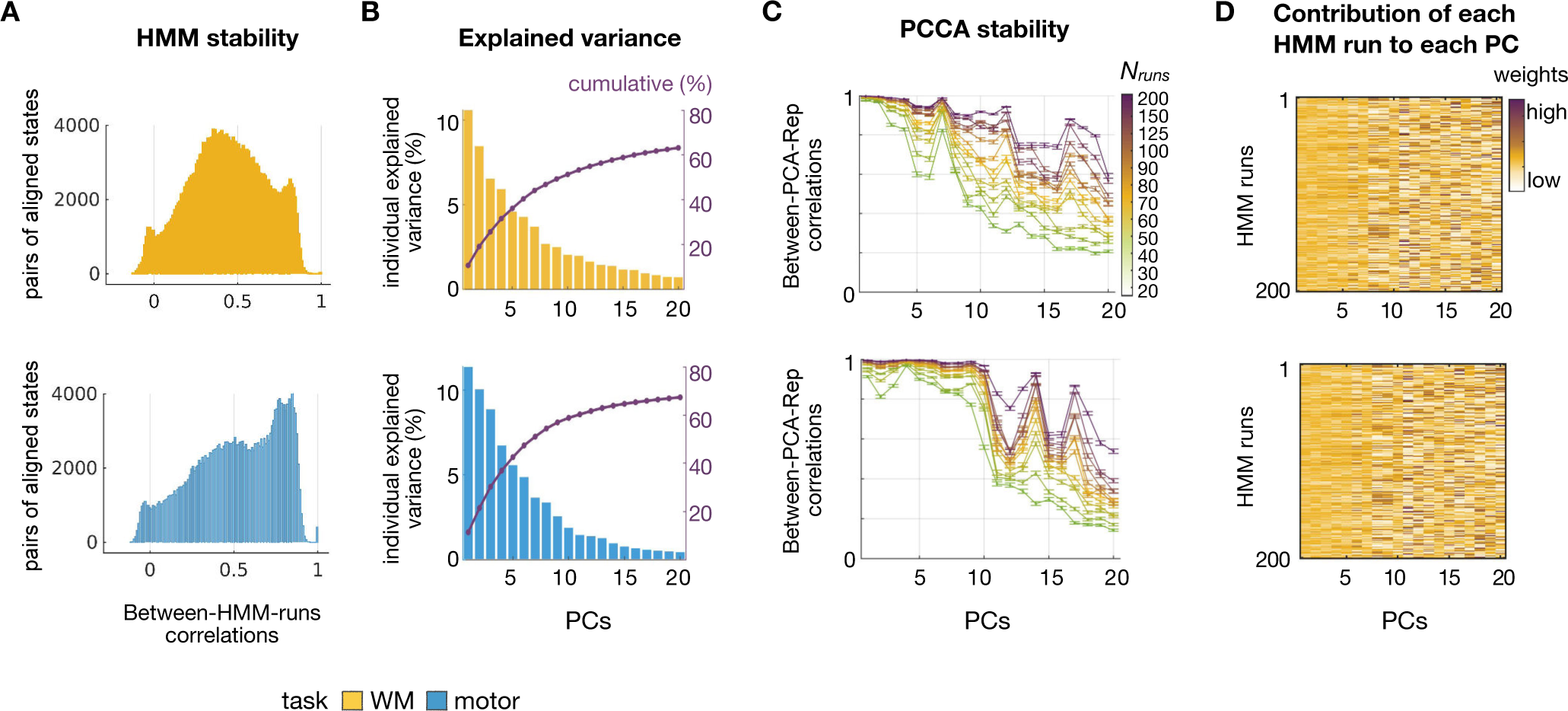
Time-varying FC trajectories in tasks span multiple temporal dimensions that can be reliably measured. For WM (top) and motor (bottom) task conditions: **A** Histogram of the Pearson correlation coeffcients between (*K* = 12) aligned states for each pair of (200) HMM runs (19900 pairs in total). **B** Plot representing the percentage of individual (bars) and cumulative (line) explained variance by the top 20 PCs that resulted from applying PCA to the K = 12 states from 200 HMM runs (i.e., 12 × 200 = 2400 states). **C** Pearson correlation coefficient (y-axis) of the 20 PCs (x-axis) between each pair of 30 PCCA repetitions, for several numbers of HMM runs (from 20 to 200); error bars represent the standard error of the mean correlation across all pairs of PCCA repetitions. **D** Average state contribution (PCA weights) of each HMM run to the final PCCA model for the top 20 PCs.

**Figure S2.**
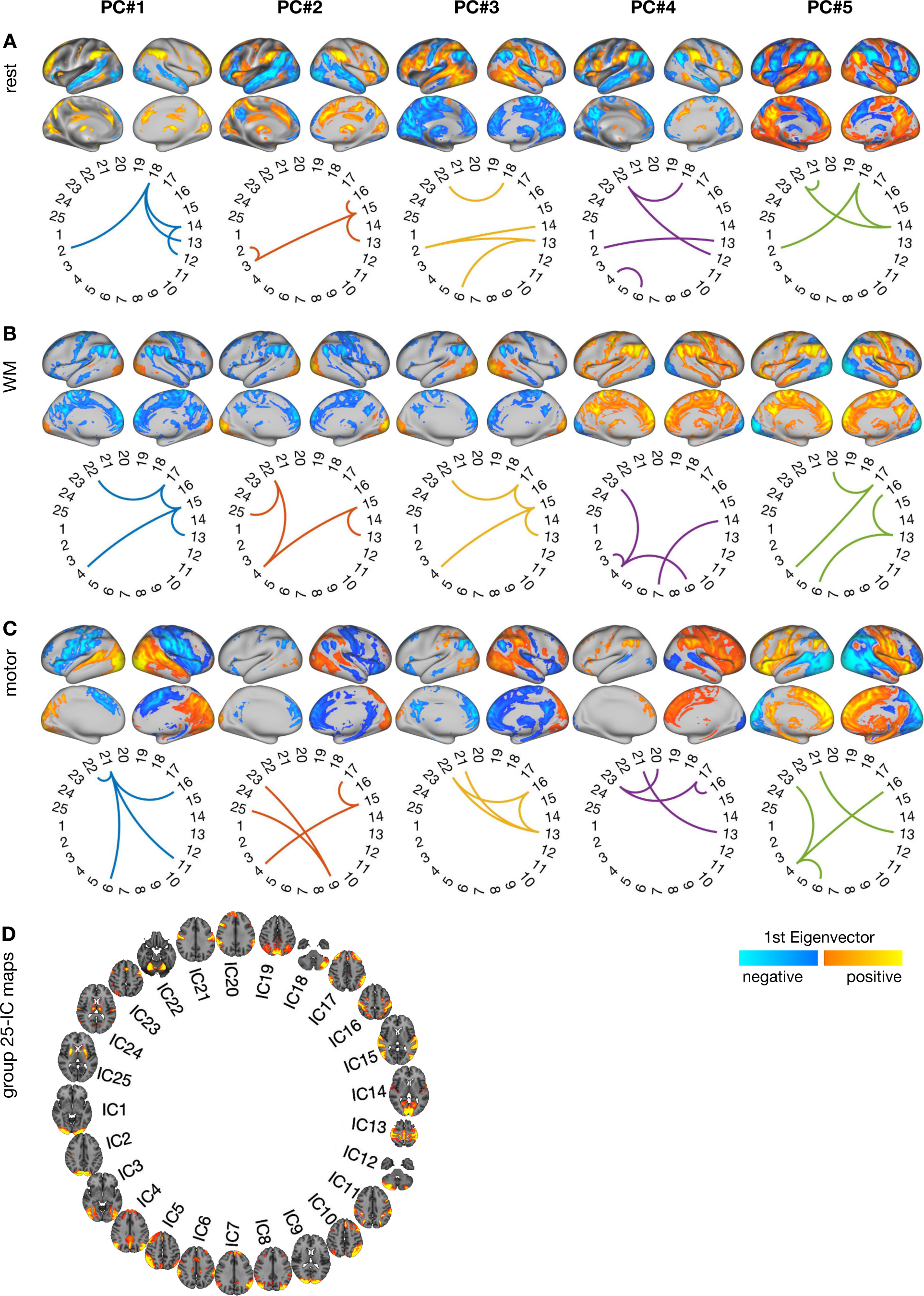
PC-specific time-varying FC patterns. (left to right) For each of the five PCs across (A) rest, (B) WM and (C) motor task conditions: (top) brain connectivity maps created using the 1^st^ Eigenvector of each PC covariance matrix. See Methods for details on the calculation of the PCs’ covariance matrices; (bottom) brain connectivity graphs of the pairwise FC across 25 ICs. D spatial maps of each IC obtained from the 25-ICA group parcellation.

**Figure S3.**
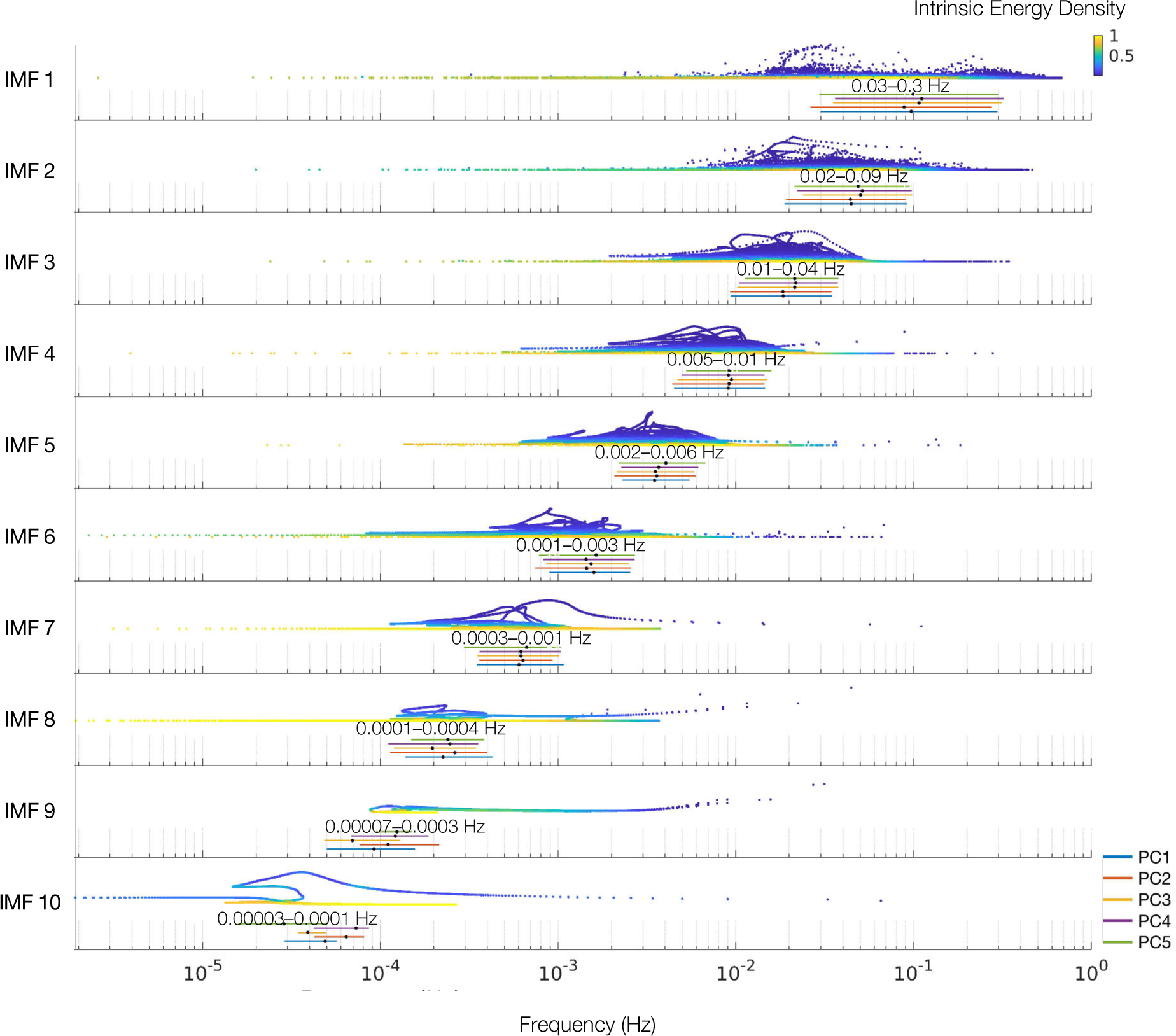
Distributions of instantaneous frequencies and instantaneous energies for each IMF, for the top five PC timeseries. Each IMF occupies a specific frequency band with little overlap between them. For each IMF, a density plot shows the instantaneous energy across frequencies, underneath, 5 horizontal lines show the median (dot) and the 90% frequency range for each PC. The instantaneous frequencies and instantaneous energies of each IMF were obtained using the Hilbert-Huang Transform.

**Table S1.** Canonical correlation analysis of the relationship between the first two moments (i.e., mean and energy) of the top 5 PC timeseries within an EMD-estimated frequency band (IMF) and the set of WM variables (i.e., cognitive load, accuracy and reaction time).

|  | IMF 1 | IMF 2 | IMF 3 | IMF 4 | IMF 5 | IMF 6 | IMF 7 | IMF 8 | IMF 9 | IMF 10 |
| --- | --- | --- | --- | --- | --- | --- | --- | --- | --- | --- |
| *Loadings* |  |  |  |  |  |  |  |  |  |  |
| SET 1 |  |  |  |  |  |  |  |  |  |  |
| mean PC1 | -0.34 | -0.2 | -0.32 | -0.34 | -0.34 | -0.1 | -0.54 | -0.68 | 0.36 | -0.13 |
| mean PC2 | -0.35 | -0.76 | -0.6 | 0.12 | 0.32 | 0.77 | -0.47 | 0.29 | -0.42 | 0.88 |
| mean PC3 | -0.13 | -0.57 | -0.49 | 0.2 | -0.067 | 0.33 | 0.2 | -0.36 | -0.53 | 1.1 |
| mean PC4 | -0.24 | -0.042 | -0.62 | 0.89 | 0.58 | 0.36 | 0.43 | 0.58 | 0.12 | 0.62 |
| mean PC5 | 0.84 | 0.25 | -0.011 | 0.28 | 0.75 | -0.45 | 0.055 | -0.27 | 0.059 | 1 |
| SET 2 |  |  |  |  |  |  |  |  |  |  |
| Load | 0.85 | -0.15 | 0.2 | 0.43 | 0.44 | 0.3 | -0.034 | 0.56 | 0.27 | -0.5 |
| ACC | 0.54 | -0.65 | -0.75 | 0.89 | 0.87 | 0.6 | 1.1 | -0.051 | -0.92 | 0.087 |
| RT | 0.28 | -0.98 | 0.31 | -0.3 | -0.34 | -0.7 | 0.62 | -1.1 | -0.93 | 1.2 |
| *r* | 0.073 | 0.073 | 0.065 | 0.069 | 0.094 | 0.076 | 0.16 | 0.19 | 0.17 | 0.18 |
| *p-Perm* | 0.15 | 0.06 | 0.0001 | 0.0001 | 0.0004 | 0.0007 | 0.93 | 0.96 | 0.66 | 0.28 |
| *p-FDR* | 0.31 | 0.13 | 0.0005 | 0.0005 | 0.0013 | 0.002 | 0.96 | 0.96 | 0.9 | 0.46 |
| *Loadings* |  |  |  |  |  |  |  |  |  |  |
| SET 1 |  |  |  |  |  |  |  |  |  |  |
| energy PC1 | 0.62 | -0.051 | -0.33 | 0.088 | 0.58 | -0.83 | 0.44 | 0.049 | 0.71 | 1 |
| energy PC2 | -0.61 | -0.92 | -0.21 | -0.59 | 0.3 | -0.36 | 0.11 | -0.39 | -0.47 | 0.35 |
| energy PC3 | -0.13 | 0.36 | -0.24 | 0.25 | 0.036 | 0.21 | -0.57 | -0.52 | 0.17 | -0.6 |
| energy PC4 | 0.38 | -0.21 | 0.56 | -0.69 | 0.21 | 0.29 | 0.59 | 0.64 | 0.27 | 0.42 |
| energy PC5 | -0.37 | 0.064 | 0.61 | 0.28 | -0.73 | -0.011 | -0.047 | -0.62 | 0.6 | 0.72 |
| SET 2 |  |  |  |  |  |  |  |  |  |  |
| Load | 0.65 | -0.028 | 0.036 | -0.48 | 0.94 | -0.064 | 0.28 | -0.38 | 0.49 | 0.5 |
| ACC | -0.75 | -0.33 | 0.95 | -1 | -0.1 | 1 | -0.83 | -0.59 | 0.34 | 0.012 |
| RT | -0.18 | 0.84 | 0.83 | -0.22 | 0.07 | 0.78 | -1 | 0.73 | -0.94 | -1.1 |
| *r* | 0.069 | 0.085 | 0.098 | 0.077 | 0.098 | 0.11 | 0.16 | 0.15 | 0.22 | 0.12 |
| *p-Perm* | 0.0002 | 0.0001 | 0.0001 | 0.021 | 0.73 | 0.81 | 0.7 | 0.81 | 0.23 | 0.76 |
| *p-FDR* | 0.0008 | 0.0005 | 0.0005 | 0.052 | 0.9 | 0.9 | 0.9 | 0.9 | 0.42 | 0.9 |

ACC: accuracy; *p-FDR*: false discovery rate adjusted *p-value* (for 2 moments x 10 IMFs = 20 tests); *p-Perm*: *p-value* of a 10000 permutation test (permutations were performed within each subject across blocks and sessions); *r*: canonical correlation coefficient; RT: reaction time; WM: working memory

**Table S2.** Canonical correlation analysis of the relationship between the first two moments (i.e., mean and energy) of the top 5 PC timeseries within an EMD-estimated frequency band (IMF) and the set of WM variables (i.e., accuracy and reaction time) for 2-back blocks (i.e., high cognitive load).

|  | IMF 1 | IMF 2 | IMF 3 | IMF 4 | IMF 5 | IMF 6 | IMF 7 | IMF 8 | IMF 9 | IMF 10 |
| --- | --- | --- | --- | --- | --- | --- | --- | --- | --- | --- |
| *Loadings* |  |  |  |  |  |  |  |  |  |  |
| SET 1 |  |  |  |  |  |  |  |  |  |  |
| mean PC1 | -0.89 | 0.13 | 0.22 | -0.44 | 0.25 | -0.8 | 0.57 | -0.37 | 0.2 | 0.27 |
| mean PC2 | -0.5 | 0.075 | -0.52 | -0.11 | 0.3 | 0.34 | -0.32 | 0.46 | -0.57 | -0.25 |
| mean PC3 | 0.086 | 0.44 | 0.41 | 0.59 | -0.57 | 0.41 | 0.34 | -0.78 | -0.36 | 0.32 |
| mean PC4 | -0.098 | 0.34 | -0.74 | -0.3 | 0.31 | 0.17 | -0.43 | 0.11 | 0.82 | -0.14 |
| mean PC5 | -0.048 | 0.85 | 0.21 | 0.29 | -0.18 | -0.037 | 0.7 | -0.17 | -0.38 | 0.9 |
| SET 2 |  |  |  |  |  |  |  |  |  |  |
| 2back ACC | 0.99 | 1 | -0.15 | -1 | 1 | 0.099 | -0.059 | 0.26 | 0.35 | 0.54 |
| 2back RT | -0.057 | 0.026 | -1 | -0.15 | 0.073 | -0.98 | 0.99 | -0.92 | 1 | 0.94 |
| *r* | 0.094 | 0.13 | 0.12 | 0.17 | 0.22 | 0.18 | 0.18 | 0.21 | 0.25 | 0.12 |
| *p-Perm* | 0.65 | 0.85 | 0.087 | 0.76 | 0.91 | 0.39 | 0.34 | 0.064 | 0.19 | 0.28 |
| *p-FDR* | 0.87 | 0.97 | 0.58 | 0.95 | 0.97 | 0.71 | 0.67 | 0.58 | 0.67 | 0.67 |
| *Loadings* |  |  |  |  |  |  |  |  |  |  |
| SET 1 |  |  |  |  |  |  |  |  |  |  |
| energy PC1 | -0.25 | 0.56 | -0.2 | 0.21 | 0.088 | -0.93 | 0.34 | 0.025 | 0.52 | -0.69 |
| energy PC2 | -0.82 | 0.57 | -0.32 | 0.27 | -0.79 | -0.008 | 0.23 | 0.12 | -0.6 | 0.68 |
| energy PC3 | 0.34 | 0.2 | -0.015 | -0.32 | -0.28 | 0.083 | 0.088 | 0.95 | -0.13 | 0.28 |
| energy PC4 | -0.18 | 0.3 | -0.33 | -0.9 | 0.18 | -0.17 | 0.85 | 0.27 | 0.93 | 0.32 |
| energy PC5 | -0.39 | -0.25 | 0.82 | -0.074 | 0.53 | 0.3 | -0.052 | -0.17 | 0.51 | -0.59 |
| SET 2 |  |  |  |  |  |  |  |  |  |  |
| 2back ACC | -1 | 0.96 | -1 | 0.76 | 0.65 | 0.6 | -1 | 0.69 | 0.47 | -0.12 |
| 2back RT | -0.16 | 0.48 | -0.09 | -0.53 | 0.88 | 0.91 | -0.2 | 0.85 | -0.81 | 0.97 |
| *r* | 0.091 | 0.15 | 0.18 | 0.12 | 0.069 | 0.15 | 0.24 | 0.18 | 0.27 | 0.21 |
| *p-Perm* | 0.56 | 0.65 | 0.24 | 0.44 | 0.3 | 0.33 | 0.16 | 0.93 | 0.039 | 0.97 |
| *p-FDR* | 0.86 | 0.87 | 0.67 | 0.73 | 0.67 | 0.67 | 0.67 | 0.97 | 0.58 | 0.97 |

ACC: accuracy; *p-FDR*: false discovery rate adjusted *p-value* (for 2 moments x 10 IMFs = 20 tests); *p-Perm*: *p-value* of a 10000 permutation test (permutations were performed between subjects accounting for the family structure of the HCP data); *r*: canonical correlation coefficient; RT: reaction time; WM: working memory

